# Transcriptome analysis of the *Larimichthys polyactis* under heat and cold stress

**DOI:** 10.1101/779462

**Authors:** Tianqi Chu, Feng Liu, Gaochan Qin, Wei Zhan, Mengjie Wang, Bao Lou

## Abstract

The small yellow croaker (*Larimichthys polyactis*) is an important marine economic fish that is widely distributed in the East Sea, Yellow Sea and Bohai of China. However, the wild populations of small yellow croaker are severely depleted, and there is currently a developing large-scale artificial propagation of this fish for aquaculture. However, the current variety of small yellow croaker that is cultivated is not capable to coping with large fluctuations in temperature. Therefore, it is important to understand the molecular mechanisms that are activated in response to temperature stress in the small yellow croaker. Here, we conducted transcriptomic analysis of the liver of small yellow croaker under heat and cold stress. A total of 270,844,888, 265,727,006 and 259,666,218 clean reads were generated from heat temperature group, low temperature group and control group, respectively, and comparing expression of genes in these transcriptomes,10,878 unigenes that were differential expressed were identified. Sixteen of the differentially expressed unigenes were validated by qRT-PCR. Pathway enrichment analysis identified that the ER pathway, immune signaling pathway and metabolic response pathway were affected by temperature stress. The results of this study provide a comprehensive overview of temperature stress-induced transcriptional patterns in liver tissues of the small yellow croaker. In addition, these results can guide future molecular studies of heat and cold stress response in this species for improving the stock used for aquaculture.

## 1. Introduction

Water temperature is considered to be the ‘abiotic master factor’ for fish (Brett J R 1971), impacting biological processes in aquatic animals, including development, growth, reproduction, metabolism, behavior, and geographic distribution (Donaldson M R et al. 2008; Somero G N 2010). Although fish can adapt to variations in water temperature through physiological plasticity or micro-evolution, severe diseases or death occurs when they are exposed to temperatures exceeding their thermal tolerance capability (Whitehead A et al. 2011; Céline Bellard et al. 2012). Due to the intensification of global warming, extremely low winter temperatures and high summer temperatures have led to mass mortality in farmed fish and damaging the economic profit of aquaculture farms. This has negatively affected the economic development of several countries where the economy is dependent on aquaculture, including China, Israel, and South America (Ibarz A et al. 2010). Therefore, investigating the mechanisms underlying temperature adaptation and tolerance in fish species that are used for aquaculture is very important.

The small yellow croaker (*Larimichthys polyactis*) belongs is a warm-temperate near-bottom migratory fish. It is widely distributed in the East Sea, Yellow Sea and Bohai of China (Li Z et al. 2011). The small yellow croaker is nutritious and its meat is of high quality, making it an important marine economic fish. However, due to overfishing and deterioration of the environment in its native habitat, the wild resources of the small yellow croaker are severely depleted (Chen WM and Cheng QQ 2013). Artificial breeding techniques of the small yellow croaker have been developed recently, laying the foundation for large-scale artificial propagation of this species (Liu F et al. 2019). The wild population of small yellow croaker migrates based on water temperature, remaining in areas that are suitable for their survival during the different seasons (Johnson J A and Kelsch S W 1998). However, fish in aquaculture cannot move beyond the cultured space and area, therefore they cannot migrate to higher or lower temperature to survive changes in temperature. The current method to combat this problem is through adjusting water temperature using thermostat, but this method increases production costs. Therefore, identifying economic methods to increase survival of small yellow croaker under changing temperatures is important for sustainable aquaculture.

Transcriptomics approaches are robust and reliable for identifying genetic pathways that are important for response to various conditions, facilitating the exploration of global gene expression changes caused by abiotic stress. Multiple studies have identified gene expression responses elicited by temperature stress in various fish species, such as rainbow trout (*Oncorhynchus mykiss*) (Li Y et al. 2017), zebrafish (*Danio rerio*) (Long Y et al. 2013), grass carp (*Ctenopharyngodon idellus*) (Yang Y et al. 2016), yellow drum (*Nibea albiflora*) (Dongdong X et al. 2018), channel catfish (*Ictalurus punctatus*) (Ju Z et al. 2002), gilthead sea bream (*Sparus aurata*) (Mininni A N et al. 2014), and large yellow croaker (*Larimichthy crocea*) (Qian B and Xue L 2016).

However, transcriptomic profile of *L. polyactis* under heat and cold temperatures stress remains to be elucidated. In the present study, RNA-Seq analysis was conducted to evaluate the effects of temperature stress on *L. polyactis*. The liver is involved in lipid, glucose and protein metabolism of fish (Tan P et al. 2017), but also regulates stress response. Therefore, we focused on the liver tissues for transcriptome analysis. Investigating the molecular mechanisms underlying temperature stress response in *L. polyactis* will contribute to understanding its adaptation to changes in temperature and promote the development of temperature-tolerant breeds for this species.

## 2. Materials and methods

### 2.1 Experimental design and sampling

One hundred and eighty small yellow croakers (55.0 ± 0.6 g) were obtained from the Marine Fishery Institute of Zhejiang Province (Xishan Island, Zhoushan, China). The fish were randomly divided into three groups with three tanks per group: heat temperature group (HTS), low temperature group (LTS) and control group (CT). Fish were cultured in aerated water tanks (0.5 m^3^) with 20 fish in each tank, and the tank had a flow through sea water supply maintained at 20 °C. After 14 days of acclimation, fish were placed under heat and cold stress by altering the temperature of the water in the tank at a constant rate of 2 °C per 1 h. In the heat temperature group, temperature was increased from 20 °C to 32 °C, and the cold temperature group was decreased from 20 °C to 6 °C. Liver tissues were sampled from five fish per group at 6 h, and samples were immediately flash-frozen in liquid nitrogen, and then stored at −80 °C for further analysis. All the procedures performed on animals were approved by the Guidelines for the Care and Use of Laboratory Animals in China.

### 2.2 RNA isolation, library preparation and sequencing

Total RNA of liver tissues was extracted using Trizol reagent (Invitrogen, CA, USA) in accordance with the manufacturer’s procedure. The total RNA quantity and purity were determined by a Bioanalyzer 2100 and RNA 1000 Nano LabChip Kit (Agilent, CA, USA) and samples with an RNA integrity number (RIN) above 7.0. Poly(A) RNA was purified from total RNA (5μg) using poly-T oligo-attached magnetic beads with the two rounds of purification. The mRNA was then fragmented into small pieces using divalent cations under elevated temperature. The cleaved RNA fragments were reverse-transcribed to create the final cDNA library for the mRNA-Seq sample preparation kit (Illumina, San Diego, USA), with an average of 300 bp (±50 bp) insert size for the paired end libraries. The paired-end sequencing was performed on Illumina Hiseq4000 (LC Sceiences, USA) following the vendor’s recommended protocol and the raw data files were deposited to NCBI’s Sequence Read Archive (SRA). The accession numbers are SRR10082705 (CT1), SRR10082704 (CT2), SRR10082698 (CT3), SRR10082697 (CT4), SRR10082696 (CT5), SRR10082703 (LTS1), SRR10082702 (LTS2), SRR10082701 (LTS3), SRR10082700 (LTS4), SRR10082699 (LTS5), SRR10082695 (HTS1), SRR10082694 (HTS2), SRR10082693 (HTS3), SRR10082692 (HTS4), SRR10082691 (HTS5).

### 2.3 *De novo* assembly, unigene annotation and functional classification

Cutadapt (Martin M 2011) and in house perl scripts were used to remove the reads that contained adaptor contamination, low quality bases and undetermined bases. Sequence quality of the clean reads were verified by FastQC (http://www.bioinformatics.babraham.ac.uk/projects/fastqc/) including the Q20, Q30 and GC-content. All downstream analyses were based on clean reads, which were of high quality. *De novo* assembly of the transcriptome was performed with Trinity 2.4.0 (Grabherr et al. 2011). Trinity groups transcripts into clusters based on shared sequence content.

All assembled unigenes were aligned against the non-redundant (Nr) protein (http://www.ncbi.nlm.nih.gov/), Gene ontology (GO) (http://www.geneontology.org), SwissProt (http://www.expasy.ch/sprot/), Kyoto Encyclopedia of Genes and Genomes (KEGG) (http://www.genome.jp/kegg/) and eggNOG (http://eggnogdb.embl.de/) databases using DIAMOND (Buchfink B et al. 2014) with a threshold of E-value<0.00001. The assembled transcriptome was used as the reference for the differential expression analysis.

### 2.4 Differential expression analysis

The expression levels of a unigene was calculated using the Transcripts per million (TPM) method (Günter P. Wagner et al. 2012) and unigenes that were significantly up-regulated or down-regulated were identified for three comparisons: HTS–LTS, HTS–CT, and LTS–CT. The differentially expressed unigenes were selected with |log2 (fold change) | >1, with statistical significance (*P*< 0.05) using R package edgeR (Robinson M D et al. 2010) and the False Discovery Rate (FDR) set to < 0.001. According to these criteria, smaller FDR and larger ratio indicates a larger difference of the expression between the compared groups. Finally, the different expression unigenes (DEUs) that were identified were used to analyze differences in Gene ontology (GO) functional categories and KEGG pathways. GO function analysis (including GO functional classification annotation and GO functional enrichment analysis) for DEUs were performed by mapping all DEUs to each term of the GO database (http://www.geneontology.org/) and calculating the gene numbers each GO term had. All of raw data about transcriptome were submitted to the NCBI GEO database.

### 2.5 Validation using quantitative real-time PCR

To examine the reliability of the RNA-Seq results, sixteen DEUs were selected based on their potential functional important for validation using quantitative real-time RT-PCR (qRT-PCR). The housekeeping gene *β-actin* was used as the reference gene. Suitable primers (Table 1) were designed using Primer Express 6.0 and synthesized by GENEWIZ (Suzhou, China) Co., Ltd. PCRs for each sample were performed as triplicates with the SYBR Green dye (TaKaRa, Dalian, China) and StepOnePlus™ Real-Time PCR System according to the manufacturer’s protocol. RNA samples that were used for the RNA-seq experiment were selected for this experiment. The qRT-PCR conditions were as follows: 95 °C for 30 s, 40 cycles at 95 °C for 5 s and 60 °C for 34 s and dissociation curve analysis was carried out to determine the target specificity. The relative expression ratio of the target genes versus *β-actin* was calculated using 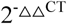 method (Schmittgen T D and Livak K J 2008). Relative mRNA expression levels were statistically analyzed using SPSS 21 software.

**Table 1:**
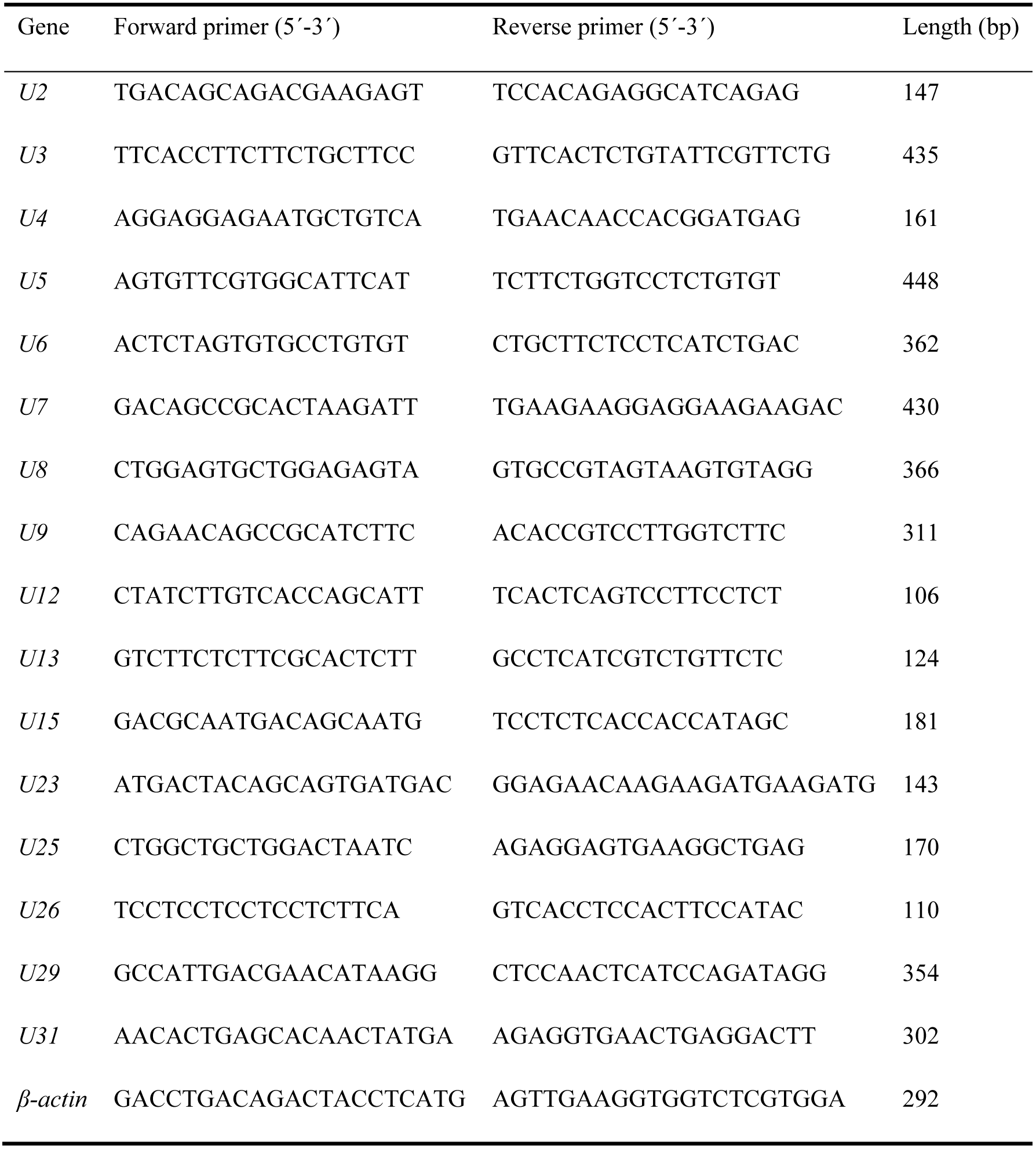
Primers used for qRT-PCR verification of differently expressed genes.

## 3. Results

### 3.1 Transcriptome assembly and annotation

To identify genes that are differentially expressed under temperature stress, transcriptomes were generated from liver samples of animals placed in the heat temperature group (HTS), low temperature group (LTS) and control group. HTS was exposed to 32 °C and LTS was exposed to 6 °C, while the control group was maintained at 20 °C. In total, 276,893,528, 271,842,088, and 265,003,108 raw reads were obtained from HTS, LTS and CT, respectively. A total of 270,844,888, 265,727,006 and 259,666,218 clean reads were generated by filtering the raw reads from HTS, LTS and CT, respectively. We predicted 86,584 transcripts from the clean reads and N50 length was 1,773 bp (Table 2). The length distribution of the transcriptome libraries is shown in Fig. 1.

**Table 2:**
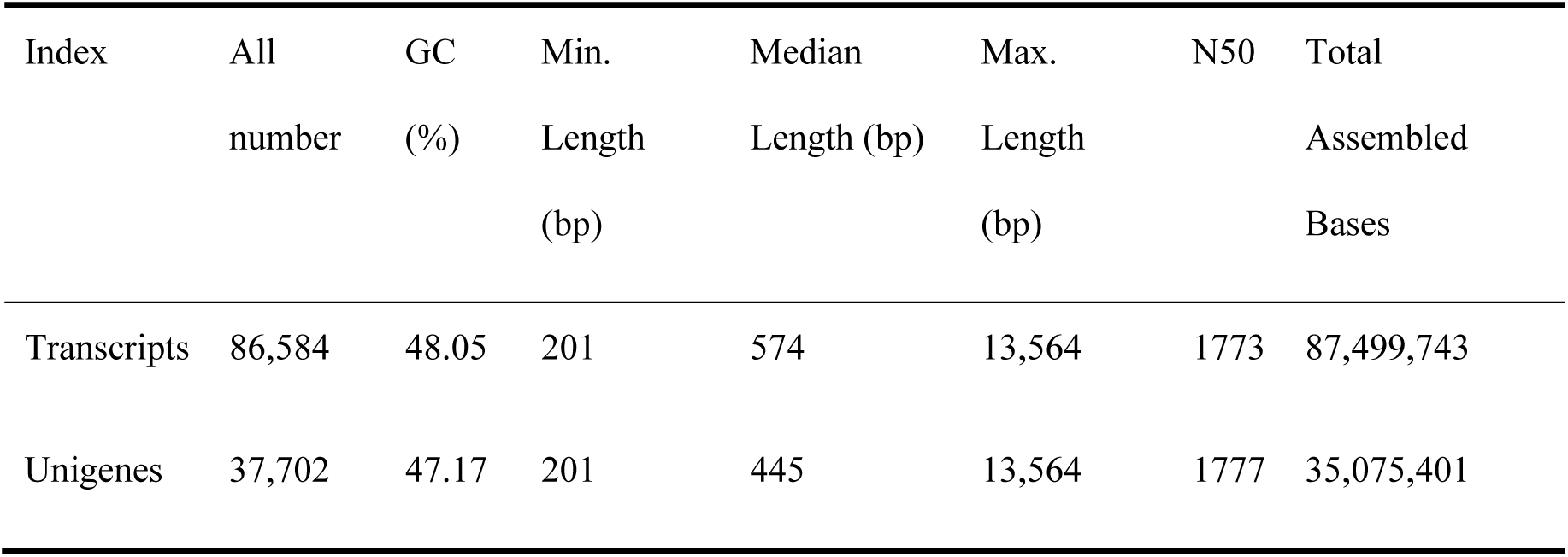
Overview of assembly results.

**Fig. 1:**
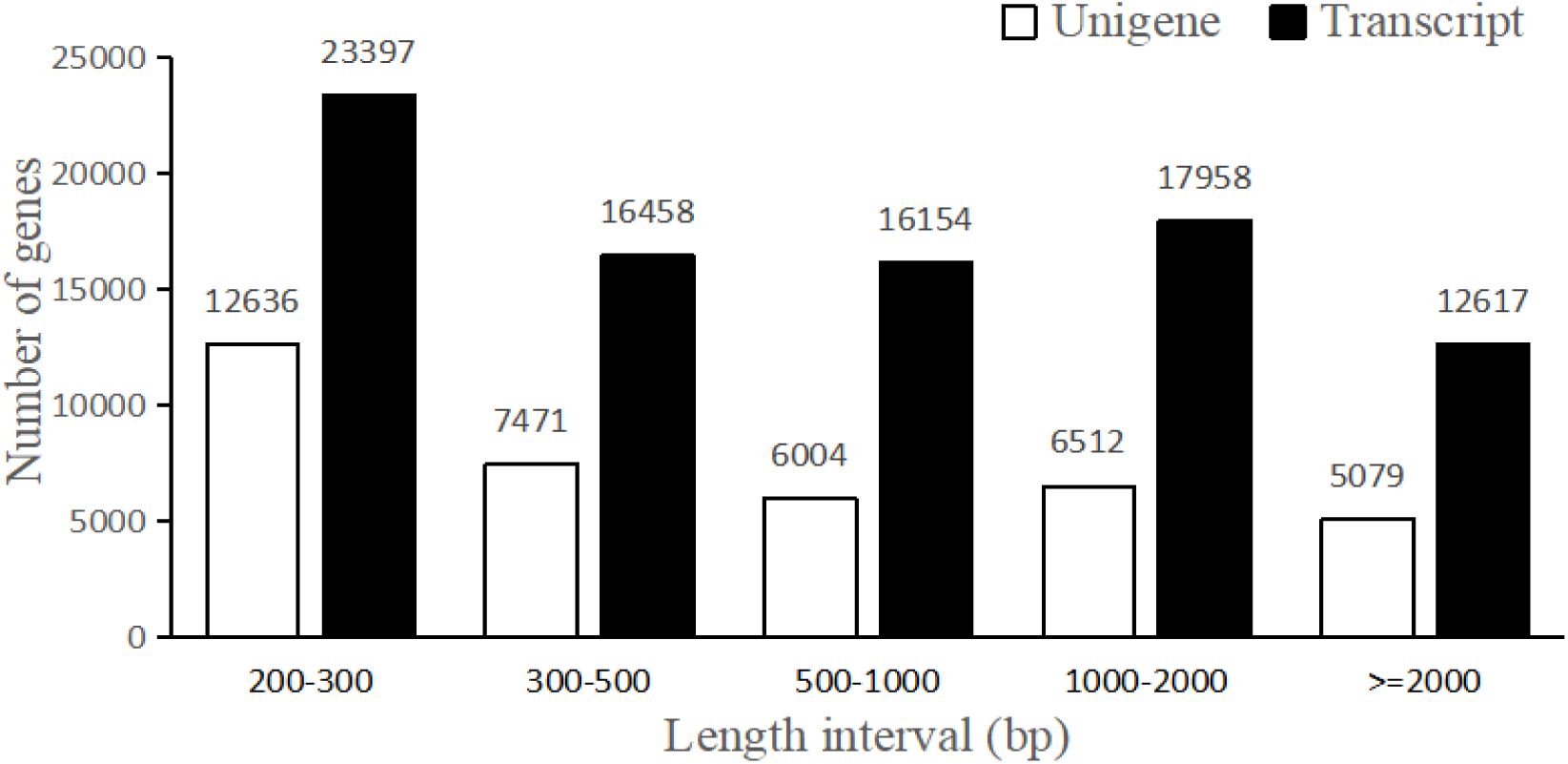
Length distributions of the transcripts and unigenes.

The value of Pearson’s correlation coefficients (R) for sample expression was 0.6-0.97 for CT1, CT2, CT3, CT4 and CT5, 0.69-0.977 for HTS1 HTS2 HTS3 HTS4 and HTS5, and 0.741-1 for LTS1 LTS2 LTS3 LTS4 and LTS5 (Fig. 2). This highlighted that there was strong correlation of gene expression patterns in the replicate liver transcriptomes, demonstrating that collection methods were accurate. Unigenes were functionally annotated using NCBI_nr, GO, KEGG, Pfam, Swiss-Prot and eggNOG (Table 3).

**Table 3:** Summary statistics of transcriptome annotation.

| Database | Number | Ratio (%) |
| --- | --- | --- |
| All | 37,702 | 100.00 |
| GO | 14,976 | 39.72 |
| KEGG | 9,587 | 25.43 |
| Pfam | 13,179 | 34.96 |
| eggNOG | 16,580 | 43.98 |
| NR | 18,272 | 48.46 |

**Fig. 2:**
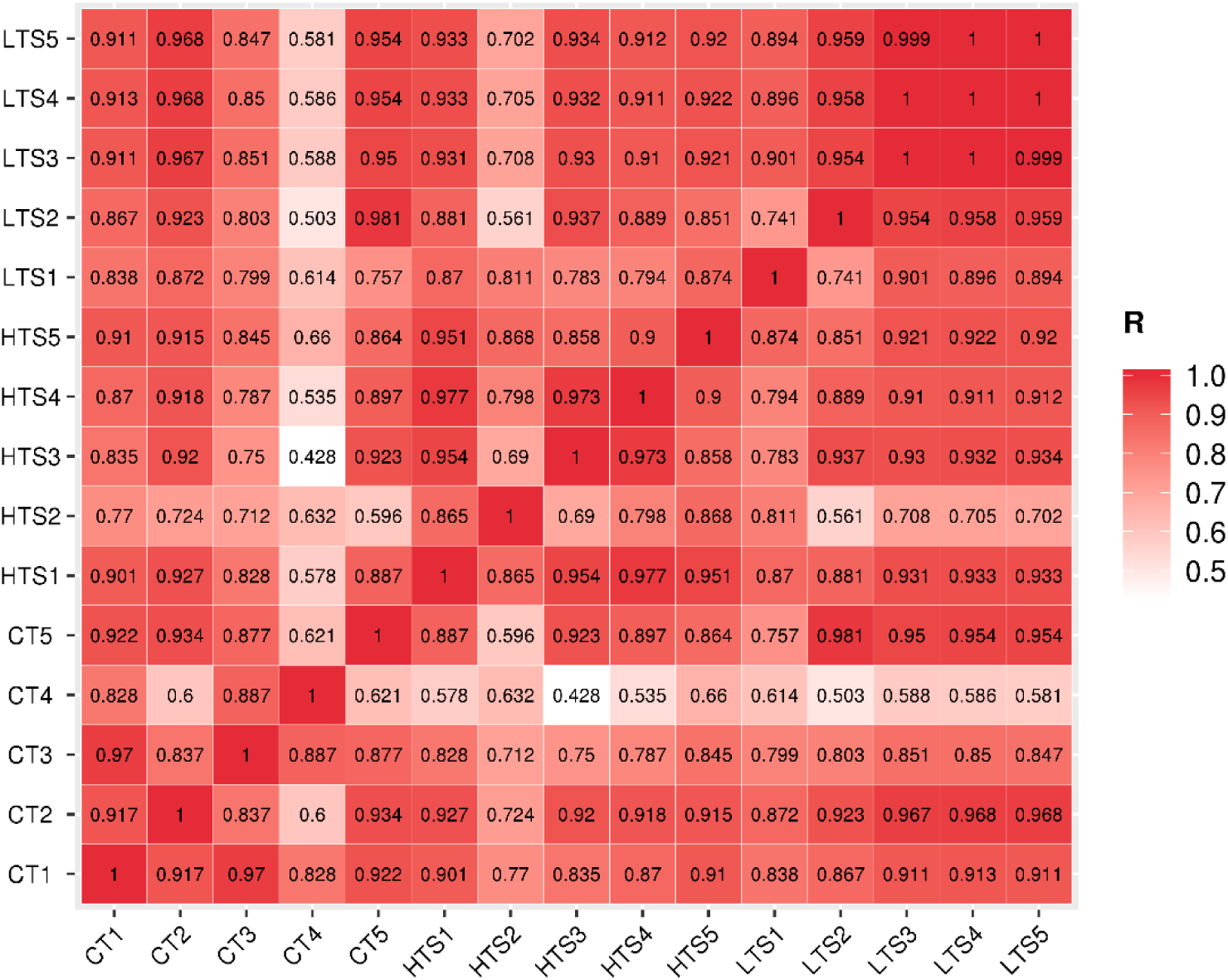
Pearson correlation between samples.

### 3.2 Differential expression analysis

We constructed a heat map to identify patterns of unigene expression across the three treatment groups (Fig.3). Of these, 1,843 unigenes were expressed only in HTS-CT, 774 unigenes were expressed only in LTS-CT, and 1,248 unigenes were expressed only in LTS-HTS, and expression of 664 unigenes were shared across all groups. Compared to CT, 4,033 unigenes were upregulated and 3530 were downregulated in HTS, 1,997 unigenes were upregulated and 2,528 were downregulated in LTS, and 4,053 unigenes were upregulated and 2,774 were downregulated in both HTS and LTS (Fig.4).

**Fig. 3:**
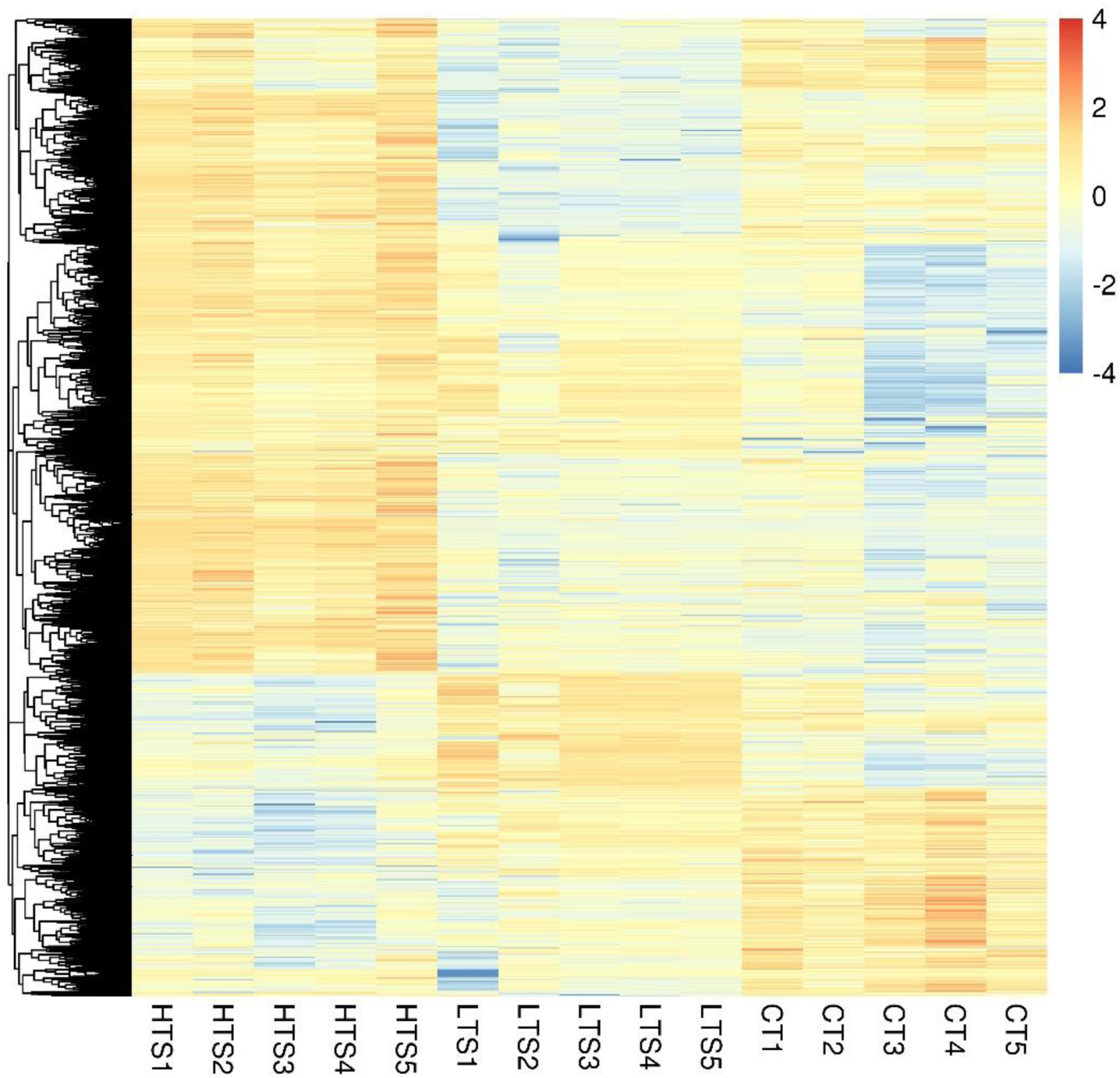
Heat map of the differentially expressed unigenes.

**Fig. 4:**
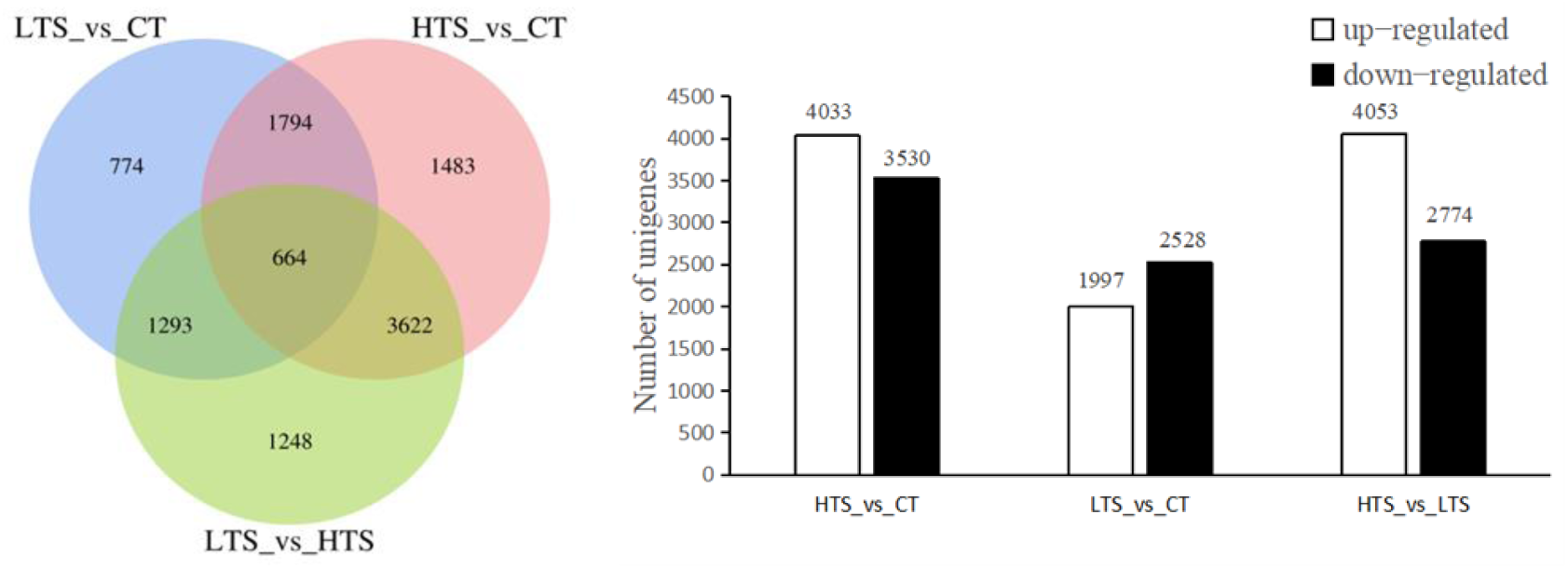
Venn diagrams of differential expressed unigenes and differentially expressed unigenes in different groups.

### 3.3 GO enrichment and KEGG pathway analysis

To evaluate the biological and functional implications of the differentially expressed unigenes (DEUs) under heat and cold stress, GO and pathway enrichment analysis were performed using the GO and KEGG databases. There were 29 significant GO categories that were enriched in HTS, LTS and CT groups (*p* < 0.01) belonging to three categories: biological process (10 subclasses), cellular component (6 subclasses) and molecular function (13 subclasses) (Fig.5). KEGG analysis identifies the associated biological signaling pathways of unigenes (Fig. 6), and we found that 13 pathways were significant enriched under HTS, LTS and CT groups (*p* < 0.01). “Protein processing in endoplasmic reticulum (ER)” was the most enriched pathway, with 256 DEUs in this pathway. Other representative pathways including “JAK-STAT signaling pathway”, “NOD-like receptor signaling pathway”, “Retinol metabolism”, “Spliceosome”, and “Cysteine and methionine metabolism” were also identified.

**Fig. 5:**
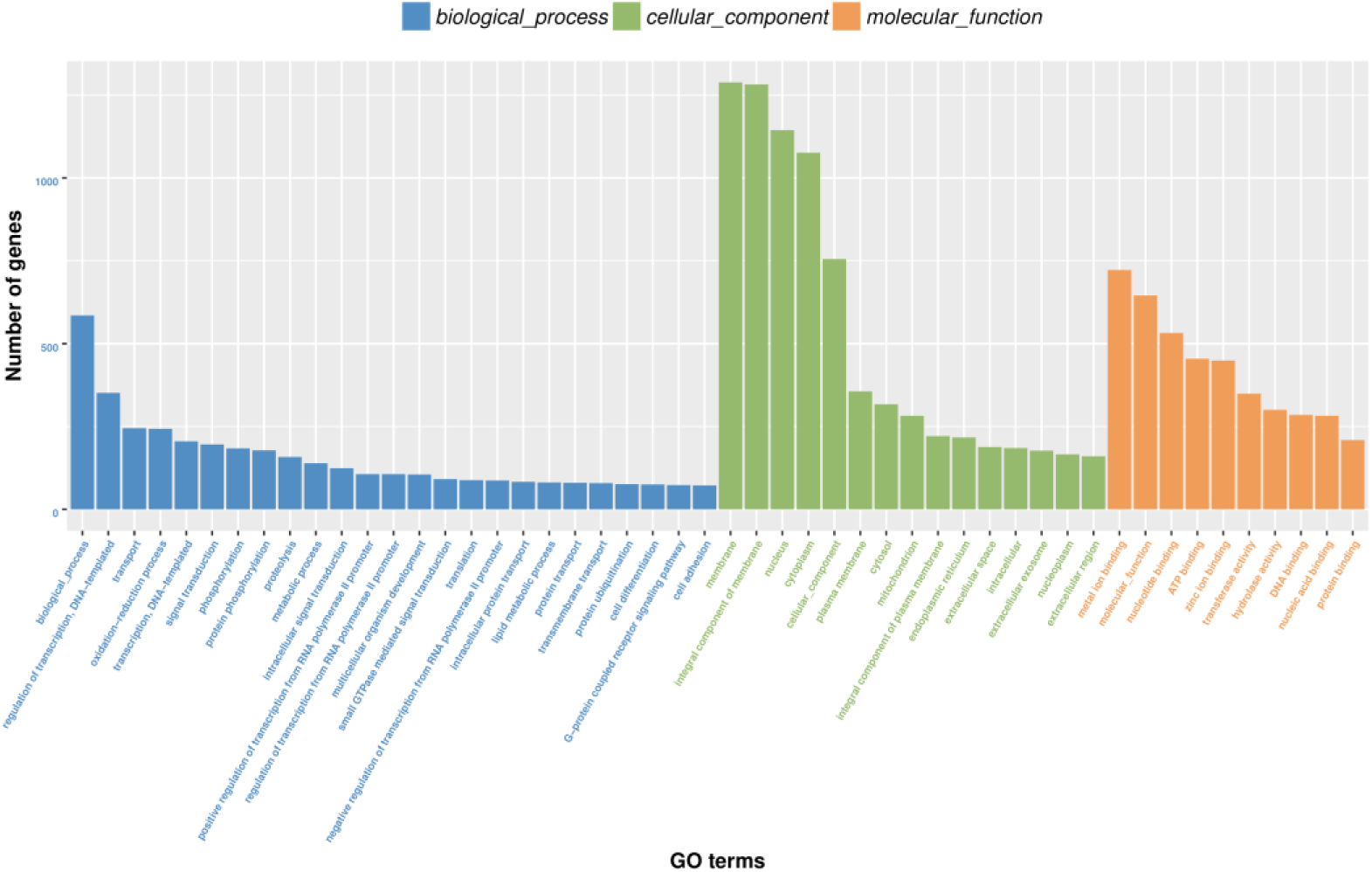
Gene ontology (GO) classifications of DEUs in HTS-LTS-CT.

**Fig. 6:**
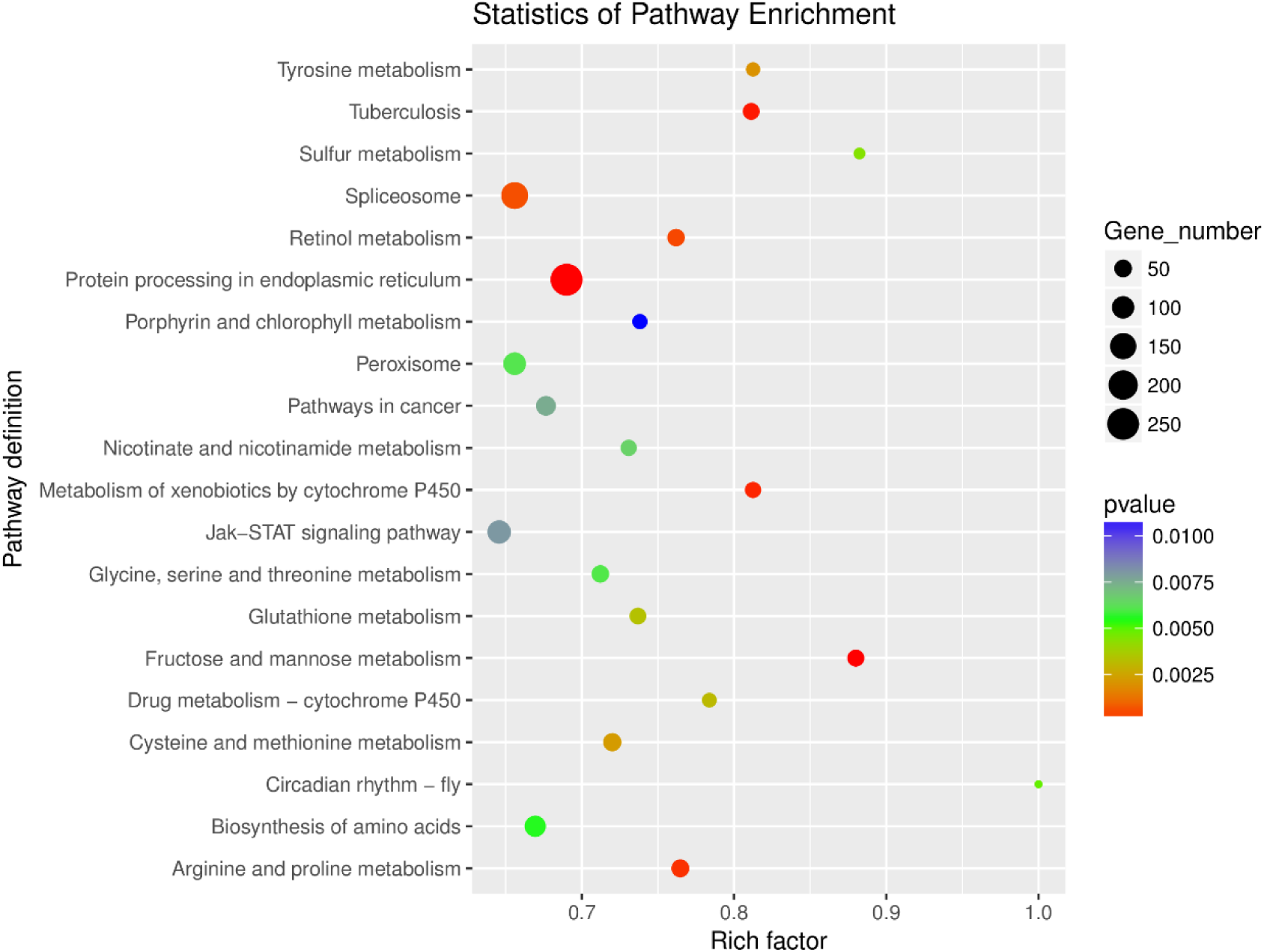
Scatterplot of enriched KEGG pathways for DEUs in HTS-LTS-CT.

### 3.4 Validation of DEUs by using qRT-PCR

Quantitative real-time PCR was performed on 16 DEUs to validate the expression patterns of the DEUs that were identified by RNA-Seq. The qRT-PCR results were significantly correlated with the RNA-Seq results (*p* < 0.01) (Table 4), and all 16 genes showed the identical up-regulated and down-regulated patterns in both qRT-PCR and RNA-Seq experiments (Fig.7).

**Fig. 7:**
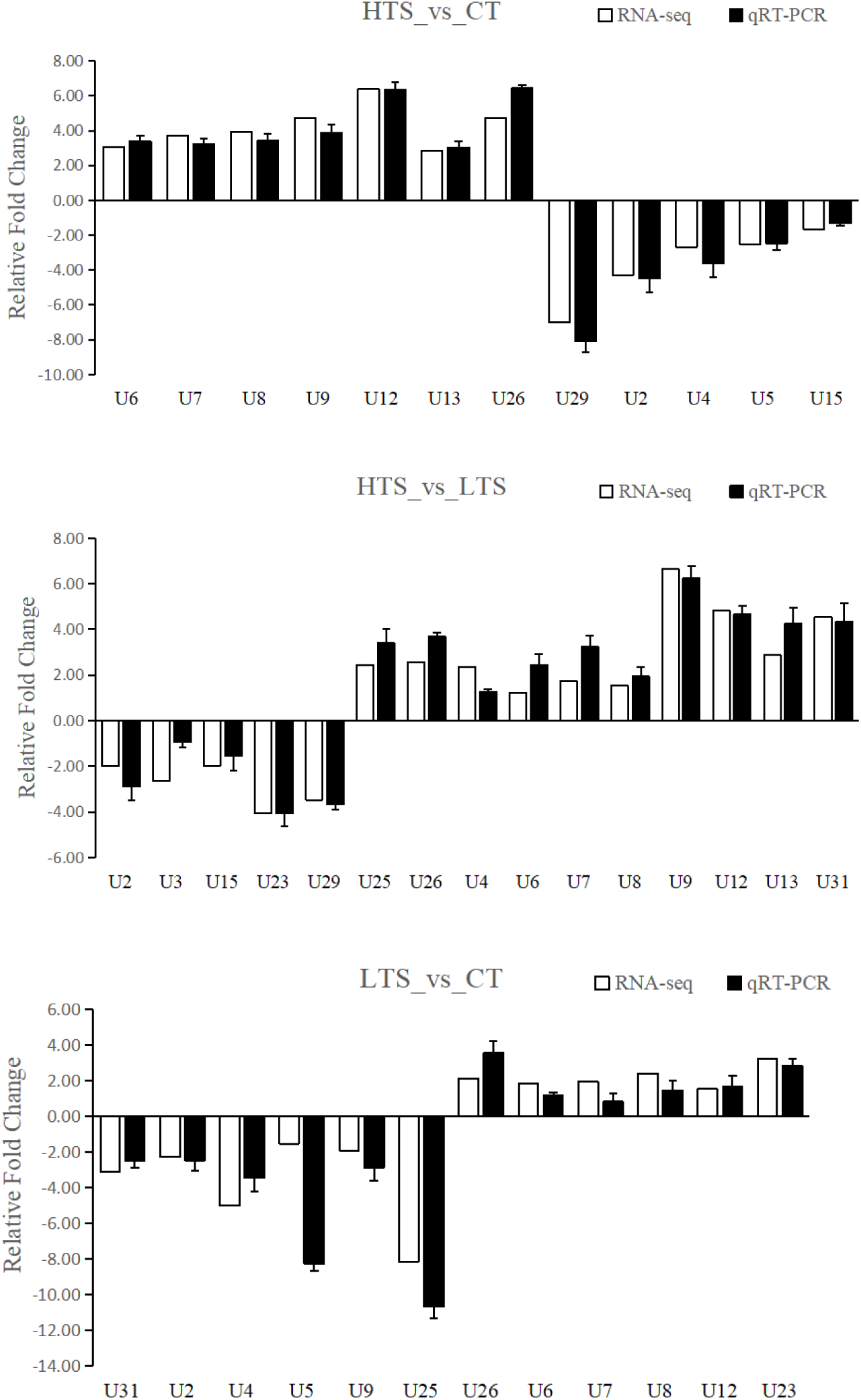
Comparison of the expressions of RNA-Seq and qRT-PCR results.

**Table 4:** The correlation coefficient of the results among RNA-Seq and real-time PCR.

| Group | Pearson correlation coefficient | Sig. |
| --- | --- | --- |
| HTS - CT | 0.988 | 0.000 |
| HTS - LTS | 0.960 | 0.000 |
| LTS - CT | 0.870 | 0.000 |

## 4. Discussion

### 4.1 The effect of temperature on the small yellow croaker

Water temperature plays an important role in the survival of small yellow croaker. Studying temperature tolerance is important to generate strains that are more resistant to changes in temperature. Under laboratory conditions, temperature tolerances of fish are usually measured either through methods where the temperature is dynamic (i.e., critical thermal methodology, CTM) or static (i.e., incipient lethal temperature, ILT) (Beitinger T L and Bennett W A 2000). Temperature tolerance has been studied extensively in several fish species. For example, the CTM temperature of redhorse suckers (*Moxostoma erythrurum*), sheepshead Minnow (*Cyprinodon variegatus*) and eastern mosqyitofish (*Gambusia holbrooki*; Poeciliidae) is 35.4°C (Reash R J et al. 2000), 45.1°C (Beitinger B T L 1997) and 40.1°C (Meffe G K et al. 1995), respectively. Previous reports have suggested that the mortality rate of small yellow croaker increases at temperatures higher than 32°C, thus we used 32°C for the HTS condition in our study. For the LTS, 6°C was selected based on the ILT temperatures of *Larimichthys crocea* (Mu F S et al. 2017).

### 4.2 Difference in gene expression patterns

In the present study, the liver transcriptome was assembled and gene expression differences in the HTS, LTS, and CT groups were compared without a reference genome, as the small yellow croaker genome has not been assembled. We predicted 86,584 transcripts and functionally annotated approximately 37,702 unigenes, which was similar to other fish species, such as Atlantic Salmon (*Salmo salar*) (Micallef G et al. 2012) and Oujiang color common carp (*Cyprinus carpio* var. color) (Du J et al. 2019). N50 can be selected as a criterion to examination the quality of gene splice. Since we did not have a genome of small yellow croaker, we estimated the N50 length using genomes of large yellow croaker, which is closely related species with it. The estimated N50 length was 1,773 bp in the small yellow croaker, which was similar to the N50 length of 1,943 bp in the large yellow croaker (Shijun X et al. 2015). The N50 value indicated that our assembled transcriptome was suitable for analysis of DEUs between three experiment groups. We next identified DEUs across the different treatment conditions, and found that heat treatment led to more up-regulated genes, and cold treatment led to more down-regulation of genes. DEUs included the Heat shock proteins (HSPs) and the Relaxin Family Peptide Receptors 3 (RXFP3).

#### 4.2.1 Heat Shock Proteins family proteins

HSPs, also known as stress proteins and extrinsic chaperones, are expressed in all organisms when exposed to stress (R.J. Roberts et al. 2010). In aquatic animals, they play a fundamental role in immune response (Dong C W et al. 2006) and inhibition of apoptosis (Sandilands J et al. 2010). In the heat stress group, 24 HSPs genes were significantly up-regulated (*p*< 0.01), and *Hsp30* had the highest expression level change (10.83-fold), followed by *Hsp90aa1* (10.11-fold), *Hsp70* (9.91-fold) and *Hsp27* (hspb1, 6.65-fold). These genes have been extensively investigated in other fish species. When rainbow trout blood was heat shocked *in vitro*, both *Hsp70* and *Hsp30* mRNA levels increased significantly (Currie S et al. 2000). *Hsp90aa1* expression is associated with stress-induced cytoprotection (Di-An F et al. 2016). *Hsp90aa1* regulates the folding of proteins under stress, as well as protein trafficking, transcriptional regulation, and epigenetic regulation of gene expression (Csermely P et al. 1998; Pearl L H and Prodromou C 2006). *Hsp27* has antiapoptotic effects through its function in the mitochondria during thermotolerance (Samali A et al.2001).

Multiple HSPs were differentially expressed in the small yellow croaker exposed to stress. *Hsp70* and *Hsp90* promotes the degradation of damaged proteins and *Hsp90a* regulates immune signaling pathway to prevent the damage. Moreover, HSPs also regulate energy metabolism, lipid, and carbohydrate levels to response to changing of temperatures. Therefore, our transcriptomic analysis suggests that increases expression of HSPs protects the small yellow croaker from heat stress.

#### 4.2.2 Relaxin Family Peptide Receptors 3

Temperature stress affects the physiology of fish at multiple levels, and can affect the central nervous system and brain activity. Under temperature stress, fish display behaviors indicative of stress and avoidance, such as erratic swimming, abnormal posture, and aggregative behavior (Quigley J T and Hinch S G 2006). Spatial and temporal ambient temperature variations directly influence cellular biochemistry and thus the physiology of fish, which are ectoderms (Burg V D and E. H 2005). Neurotransmitters regulate activity in fish (Ortiz M and Lutz P L 1995) and brain monoamine neurotransmitters induce agonistic behavior and stress reactions (Winberg S and Nilsson G E 1993). Therefore, the cranial nerve response of fish is important in its response to changes in temperature.

In the DEUs analysis, we found that *RXFP3* expression changed significantly under temperature stress (*p*< 0.01), which was up-regulated in HT vs LT (6.24-fold), down-regulated in LT vs CT (6.51-fold) and up-regulated in HT vs CT (1.63-fold). *RXFP3* regulates a wide range of behaviors, including feeding, stress responses, arousal, and cognitive processes (Bathgate R A D et al. 2013). *RXFP3* has primarily been studied in mammals (Ganella D E et al. 2013; Kania A et al. 2014), and very few studies have investigated its function marine fish (Fiengo M et al. 2013). For this reason, it is unclear whether *RXFP3* regulates neuroendocrine response to temperature. Our findings suggest that *RXFP3* plays an important role in temperature response. The mechanisms underlying this need further investigation.

### 4.3 Pathways that are differentially expressed under temperature stress

A number of enriched pathways were found in the different treatment conditions. These included protein processing in endoplasmic reticulum pathway, immune signaling pathway and metabolic response pathway, and details are discussed below.

#### 4.3.1 Protein processing in endoplasmic reticulum pathway

Endoplasmic reticulum stress (ERS) is a vital mechanism of cellular self-defense, but strong and long-lasting ERS leads to irreversible cell damage (Qing D and Zhen Z 2009). ER is a subcellular organelle where proteins are folded with the help of lumenal chaperones. Accumulation of misfolded proteins in the ER causes ER stress and activates a signaling pathway called the unfolded protein response (UPR). Studies of UPR targets engaged in endoplasmic reticulum-associated protein degradation (ERAD) reveal an intimate coordination between efficient ERAD requires an intact UPR, and inducing UPR increases ERAD capacity. Conversely, loss of ERAD leads to constitutive UPR induction (Wodicka L 2000). It is well established that the number of correctly folded protein significantly increases under heat stress. There are three ER transmembrane proteins in the small yellow croaker: endoplasmic reticulum kinase (PERK), active transcription factor 6 (ATF6) and IRE1, which promote cell survival by reducing the level of misfolded proteins (Lin J H et al. 2007). These expression levels of these genes were up-regulated in animals that were under heat stress. Due to the continued ERS, the ER function is severely impaired and the organelle elicits apoptotic signals, which can be identified by the up-regulation of C/EBP homologous protein (CHOP). During ERAD, both the Heat Shock Proteins family proteins (*Hsp40, Hsp70, Hsp90*) are up-regulated. HSPs, such as *Hsp40* and *Hsp70*, can interact with misfolded proteins, preventing them from forming aggregates (Fink A L 1999). In the goby (*Gillichthys mirabilis*), *Hsp70* and *Hsp90* were strongly up-regulated during heat shock and recovery from stress (Buckley and B. A 2006). In the study of Maraena whitefish (*Coregonus maraena*), unfolded protein response (the major ER stress pathway) was one of two characteristic pathways that were induced during acute and gradual heat stress (Alexander R et al. 2018). Our results are consistent with the above studies, showing that *Hsp40, Hsp70* and *Hsp90* were up-regulated under heat stress.

In lower temperature stress experiment, *PERK* was the only ER transmembrane protein kinases that were down-regulated, and the expression levels of *CHOP* was similar to the control. Protein Disulfide Isomerase (PDI) recognizes unfolded and partially folded proteins. Due to expression levels were down-regulated of PDIs family associated proteins, the function of PNGase and DUB were undermined causing the damage of ERAD function. However, up-regulation of *Hsp40* could compensate for the degradation level of ERAD ensuring the recovery and maintenance of liver function. In conclusion, ER function may be important for the temperature stress response of the small yellow croaker.

#### 4.3.2 Immune signaling pathway

Studies of immune pathways often focus on disease (Maekawa S et al. 2017). However, stress response and immune dysfunction are increasingly being linked through similar molecular pathways (Philip A M and Vijayan M M 2015). In general, temperature affects the immune system in teleosts (Bowden T J et al. 2007; Rebl A et al. 2013). Here, we also found changes in the expression of genes belonging to the Jak-STAT and NOD-like receptor signaling pathways under temperature stress.

The JAK-STAT signaling pathway is one of the ubiquitous signaling pathways in metazoans, and is involved in proliferation, differentiation, survival, apoptosis of cells, and mediates immune disorders (Song Z et al. 2012). JAK-STAT is negatively regulated by Protein Tyrosine Phosphatase (PTP), Suppressor of cytokine signaling (SOCS) and protein inhibitor of activated STAT (PIAS). TC-PTP can phosphorylate JAK1, JAK3, STAT1, STAT3, and STAT5 (Pouliot P et al. 2009), and the Kinase activity of JAKs and activation of STATs are inhibited by SOCS (Morales J K et al. 2010), and the transcription of STATs is inhibited by PIAS (Ivashkiv L B and Hu X 2004). Under heat stress, the expression levels of TC-PTP and PIAS were up-regulated, which likely promotes the expression levels of SOCS and activate anti-apoptotic programs. Under cold stress, STATs were down-regulated, which likely reduced the expression levels of SOCS.

In the NOD-like receptor signaling pathway, many genes were up-regulated after temperature experiment, including *NOD2* and signal mediators (*TRIP6, Nemo, NFKB* and *JNK*). The expression levels of *RIP2, CARD8, PSTPIP1* and *HSP90* family were up-regulated under heat stress and down-regulated in lower temperature stress. These expression changes are likely to indirectly influence the immune system. Thermal stress experiment in the Indian major carp catla (*Catla catla*) also found the activation of the NOD signaling pathway during thermal stress (Madhubanti Basu et al. 2015). Therefore, these immune signaling pathways are likely to play a role in temperature stress response in the small yellow croaker.

#### 4.3.3 Metabolic response pathway

The metabolism of fish is dependent on the environmental temperature, and changes metabolic rate is one of the most rapid cellular responses to the change of temperature (Somero G N 2010a). Lipid peroxidation and antioxidant defense are affected by temperature (Bagnyukova T V et al. 2007), altering energy metabolism, lipid, and carbohydrate levels through heat-shock proteins (Buckley and B. A 2006a; Vergauwen L et al. 2010).

In present study, many biological processes were significantly altered when the small yellow croaker were exposed to temperature stress. Metabolic response pathways, such as “Glutathione metabolism”, “Carbon metabolism”, “Cysteine and methionine metabolism”, “Arginine and proline metabolism”, “Retinol metabolism”, “Oxidative phosphorylation”, “Lipid and steroid metabolism”, “Fructose and mannose metabolism”, “Amino acid metabolism” and “Carbohydrate metabolism” were enriched in DEUs of animals exposed to temperature stress. This indicates that the regulation of metabolic processes plays a key role of temperature response of the small yellow croaker. In particular, metabolism of amino acids, carbohydrate and lipid were significantly influenced by the temperature treatment, such as glycine, arginine, fructose, glycolate and glucose (Rawles S D et al. 2012). Under heat stress, ATP-generating enzymes and *glucose-6-phosphatase* were up-regulated, suggesting that heat stress led to a rapid production of ATP. This may be due in part to the requirement for ATP for the function of molecular chaperones (Fink A L 1999a). Under cold stress, genes involved in lipid metabolism were down-regulated, repressing fatty acid synthase and ceramide kinase expression levels. Exposure to cold modifies lipid metabolism by lowering total saturated fatty acids in juvenile red drum (Craig S R et al. 1995). Combined together, metabolic response pathways may play an important role in the small yellow croaker’s response to temperature stress.

## 5. Conclusions

In conclusion, our transcriptome analysis demonstrated that temperature stress significantly altered gene expression in the small yellow croaker. A large number of DEUs were identified between animals that were exposed to control, heat and low temperatures. In addition, *RXFP3* was identified as a candidate gene that mediates temperature response. DEUs were enriched in the ER pathway, immune signaling pathway and metabolic response pathway. By identifying candidate genes and cellular pathways involved in temperature stress response, we provide important insights for future strategies to generate small yellow croaker breeds that are tolerant of temperature stress for aquaculture purposes.

## Acknowledgments

This research was supported by grants from the National Key Research and Development Program of China (No. 2018YFD0901204), and the Special Fund for the key research and development project of Zhejiang Province (No. 2017C02013).

## References

[1] Brett J R (1971) Energetic Responses of Salmon to Temperature. A Study of Some Thermal Relations in the Physiology and Freshwater Ecology of Sockeye Salmon (*Oncorhynchus nerkd*). American Zoologist 11(1): 99–113.

[2] Donaldson M R, Cooke S J, Patterson D A, et al. (2008) Cold shock and fish. Journal of Fish Biology 73(7): 1491–1530.

[3] Somero G N (2010) The physiology of climate change: How potentials for acclimatization and genetic adaptation will determine ‘winners’ and ‘losers’. Journal of Experimental Biology 213(6): 912–920.

[4] Whitehead A, Galvez F, Zhang S, et al. (2011) Functional Genomics of Physiological Plasticity and Local Adaptation in Killifish. Journal of Heredity 102(5): 499–511.

[5] Céline Bellard, Bertelsmeier C, Leadley P, et al. (2012) Impacts of climate change on the future of biodiversity. Ecology Letters 15(4): 365–377.

[6] Ibarz A, Francesc Padrós, Maria ángeles Gallardo, et al. (2010) Low-temperature challenges to gilthead sea bream culture: review of cold-induced alterations and ‘Winter Syndrome’. Reviews in Fish. Biology & Fisheries 20(4):539–556.

[7] Li Z, Shan X, Jin X, et al. (2011) Long-term variations in body length and age at maturity of the small yellow croaker (*Larimichthys polyactis* Bleeker, 1877) in the Bohai Sea and the Yellow Sea, China. Fisheries Research 110(1):67–74.

[8] Chen WM, Cheng QQ (2013) Development of thirty-five novel polymorphic microsatellite markers in *Pseudosciaena polyactis* (Perciformes: Sciaenidae) and cross-species amplification in closely related species, *Pseudosciaena crocea*. Biochemical Systematics and Ecology 47: 111–115.

[9] Liu F, Liu Y Y, Chu T Q, et al. (2019) Interspecific hybridization and genetic characterization of *Larimichthys polyactis* (♀) and L. crocea (♂). Aquaculture International 27:663–674.

[10] Johnson J A, Kelsch S W (1998) Effects of evolutionary thermal environment on temperature-preference relationships in fishes. Environmental Biology of Fishes 53(4): 447–458.

[11] Li Y, Huang J, Liu Z, et al. (2017) Transcriptome analysis provides insights into hepatic responses to moderate heat stress in the rainbow trout (*Oncorhynchus mykiss*). Gene 619: 1–9.

[12] Long Y, Song G, Yan J, et al. (2013) Transcriptomic characterization of cold acclimation in larval zebrafish. BMC Genomics 14(1): 612–612.

[13] Yang Y, Yu H, Li H, et al. (2016) Effect of high temperature on immune response of grass carp (*Ctenopharyngodon idellus*) by transcriptome analysis. Fish & Shellfish Immunology 58: 89–95.

[14] Dongdong X, Qiaochu Y, Changfeng C, et al. (2018) Transcriptional response to low temperature in the yellow drum (*Nibea albiflora*) and identification of genes related to cold stress. Comparative Biochemistry and Physiology Part D: Genomics and Proteomics 28: 80–89.

[15] Ju Z, Dunham R, Liu Z (2002) Differential gene expression in the brain of channel catfish (*Ictalurus punctatus*) in response to cold acclimation. Molecular Genetics & Genomics 268(1): 87–95.

[16] Mininni A N, Milan M, Ferraresso S, et al. (2014) Liver transcriptome analysis in gilthead sea bream upon exposure to low temperature. BMC Genomics 15(1): 765.

[17] Qian B, Xue L (2016) Liver transcriptome sequencing and de novo annotation of the large yellow croaker (*Larimichthy crocea*) under heat and cold stress. Marine Genomics 25: 95–102.

[18] Tan P, Dong X, Xu H, et al. (2017) Dietary vegetable oil suppressed non-specific immunity and liver antioxidant capacity but induced inflammatory response in Japanese sea bass (Lateolabrax japonicus). Fish & Shellfish Immunology 63: 139–146.

[19] Martin M (2011) Cutadapt removes adapter sequences from high-throughput sequencing reads. Embnet Journal 17(1).

[20] Grabherr M G, Haas B J, Yassour M, et al. (2011) Full-length transcriptome assembly from RNA-Seq data without a reference genome. Nature Biotechnology 29(7): 644–652.

[21] Buchfink B, Xie C, Huson D H (2014) Fast and sensitive protein alignment using DIAMOND. Nature Methods 12(1): 59–60.

[22] Günter P. Wagner, Kin K, Lynch V J (2012) Measurement of mRNA abundance using RNA-seq data: RPKM measure is inconsistent among samples. Theory Biosci 131(4): 281–285.

[23] Robinson M D, Mccarthy D J, Smyth G K (2010) EdgeR: a Bioconductor package for differential expression analysis of digital gene expression data. Biogeosciences 26: 139–140.

[24] Schmittgen T D, Livak K J (2008) Analyzing real-time PCR data by the comparative CT method. Nature Protocols 3(6): 1101–1108.

[25] Beitinger T L, Bennett W A (2000) Quantification of the Role of Acclimation Temperature in Temperature Tolerance of Fishes. Environmental Biology of Fishes 58(3): 277–288.

[26] Reash R J, Seegert G L, Goodfellow W L (2000) Experimentally-derived upper thermal tolerances for redhorse suckers: revised 316(A) variance conditions at two generating facilities in Ohio. Environmental Science and Policy 3(supp-S1): 191–196.

[27] Beitinger B T L (1997) Temperature Tolerance of the Sheepshead Minnow, *Cyprinodon variegatus*. Copeia (1): 77–87.

[28] Meffe G K, Weeks S C, Mulvey M, et al. (1995) Genetic differences in thermal tolerance of eastern mosqyitofish (*Gambusia holbrooki*; Poeciliidae) from ambient and thermal ponds. Canadian Journal of Fisheries and Aquatic Sciences 52(12): 2704–2711.

[29] Mu F S, Miao L, Li M Y, et al. (2017) Screening of microsatellite markers associated with cold tolerance of large yellow croaker (*Pseudosciaena crocea*). Journal of Biology 34(1): 34–38.

[30] Micallef G, Bickerdike R, Caroline Reiff, et al. (2012) Exploring the Transcriptome of Atlantic Salmon (*Salmo salar*) Skin, a Major Defense Organ. Marine Biotechnology 14(5): 559–569.

[31] Du J, Chen X, Wang J, et al. (2019) Comparative skin transcriptome of two Oujiang color common carp (Cyprinus carpio var. color) varieties. Fish Physiology and Biochemistry 45:177–185.

[32] Shijun X, Zhaofang H, Panpan W, et al. (2015) Functional Marker Detection and Analysis on a Comprehensive Transcriptome of Large Yellow Croaker by Next Generation Sequencing. PLOS ONE 10(4): 0124432.

[33] R.J. Roberts, C. Agius, C. Saliba, P. Bossier, Y.Y. Sung (2010) Heat shock proteins (chaperones) in fish and shellfish and their potential role in relation to fish health: a review. Journal of Fish Diseases 33(10):789–801.

[34] Dong C W, Zhang Y B, Zhang Q Y, et al. (2006) Differential expression of three *Paralichthys olivaceus* Hsp40 genes in responses to virus infection and heat shock. Fish & Shellfish Immunology 21(2): 146–158.

[35] Sandilands J, Drynan K, Roberts R J (2010) Preliminary studies on the enhancement of storage time of chilled milt of Atlantic salmon, *Salmo salar* L., using an extender containing the TEX-OE heat shock-stimulating factor. Aquaculture Research 41(4):568–571.

[36] Currie S, Moyes C D, Tufts B L (2000) The effects of heat shock and acclimation temperature on hsp70 and hsp30 mRNA expression in rainbow trout: In vivo and in vitro comparisons. Journal of Fish Biology 56(2):398–408.

[37] Di-An F, Jin-Rong D, Yan-Feng Z, et al. (2016) Molecular Characteristic, Protein Distribution and Potential Regulation of HSP90AA1 in the Anadromous Fish *Coilia nasus*. Genes 7(8): 1–12.

[38] Csermely P, Schnaider T, Soti C, et al. (1998) The 90-kDa Molecular Chaperone Family: Structure, Function, and Clinical Applications. A Comprehensive Review. Pharmacol Ther, 79(2): 129–168.

[39] Pearl L H, Prodromou C (2006) Structure and Mechanism of the Hsp90 Molecular Chaperone Machinery. Annual Review of Biochemistry 75(1): 271–294.

[40] Samali A, Robertson J D, Peterson E, et al. (2001) Hsp27 Protects Mitochondria of Thermotolerant Cells against Apoptotic Stimuli. Cell Stress & Chaperones 6(1): 49–58.

[41] Quigley J T, Hinch S G (2006) Effects of rapid experimental temperature increases on acute physiological stress and behaviour of stream dwelling juvenile chinook salmon. Journal of Thermal Biology 31(5): 429–441.

[42] Burg V D E. H (2005) Brain Responses to Ambient Temperature Fluctuations in Fish: Reduction of Blood Volume and Initiation of a Whole-Body Stress Response. Journal of Neurophysiology 93(5): 2849–2855.

[43] Ortiz M, Lutz P L (1995) Brain Neurotransmitter Changes Associated with Exercise and Stress in A Teleost Fish (*Sciaenops Ocellatus*). Journal of Fish Biology 46(4): 551–562.

[44] Winberg S, Nilsson G E (1993) Roles of brain monoamine neurotransmitters in agonistic behaviour and stress reactions, with particular reference to fish. Comparative Biochemistry and Physiology C Comparative Pharmacology and Toxicology 106(3): 597–614.

[45] Bathgate R A D, Oh M H Y, Jason L W J, et al. (2013) Elucidation of relaxin-3 binding interactions in the extracellular loops of RXFP3. Frontiers in Endocrinology 4(13): 1–10.

[46] Ganella D E, Ma S, Gundlach A L (2013) Relaxin-3/RXFP3 Signaling and Neuroendocrine Function – A Perspective on Extrinsic Hypothalamic Control. Frontiers in Endocrinology 4(128): 1–11.

[47] Kania A, Lewandowski M H, Anna Błasiak (2014) [Relaxin-3 and relaxin family peptide receptors--from structure to functions of a newly discovered mammalian brain system. Postepy Higieny I Medycyny Doswiadczalnej 68(242): 851.

[48] Fiengo M, Gaudio R D, Iazzetti G, et al. (2013) Developmental expression pattern of two zebrafish rxfp3 paralogue genes. Development Growth & Differentiation 55(9).

[49] Qing D, Zhen Z (2009) Role of endoplasmic reticulum stress in the pathogenesis of liver diseases. International Journal of Internal Medicine 36(11): 665–668.

[50] Wodicka L (2000) Functional and genomic analyses reveal an essential coordination between the unfolded protein response and ER-associated degradation. Cell 101(3): 249–258.

[51] Lin J H, Li H, Yasumura D, et al. (2007) IRE1 Signaling Affects Cell Fate During the Unfolded Protein Response. Science 318(5852): 944–949.

[52] Fink A L (1999) Chaperone-Mediated Protein Folding. Physiological Reviews 79(2): 425–449.

[53] Buckley, B. A (2006) The cellular response to heat stress in the goby *Gillichthys* mirabilis: a cDNA microarray and protein-level analysis. Journal of Experimental Biology 209(14): 2660–2677.

[54] Alexander R, Marieke V, Mareen N, et al. (2018) Gradual and Acute Temperature Rise Induces Crossing Endocrine, Metabolic, and Immunological Pathways in Maraena Whitefish (Coregonus maraena). Frontiers in Genetics 9(241).

[55] Maekawa S, Byadgi O, Chen Y C, et al. (2017) Transcriptome analysis of immune response against, Vibrio harveyi, infection in orange-spotted grouper (*Epinephelus coioides*). Fish & Shellfish Immunology 70: 628–637.

[56] Philip A M, Vijayan M M (2015) Stress-Immune-Growth Interactions: Cortisol Modulates Suppressors of Cytokine Signaling and JAK/STAT Pathway in Rainbow Trout Liver. PLOS ONE 10(1371).

[57] Rebl A, et al. (2013) Transcriptome Profiling of Gill Tissue in Regionally Bred and Globally Farmed Rainbow Trout Strains Reveals Different Strategies for Coping with Thermal Stress. Marine Biotechnology 15(4): 445–460.

[58] Bowden T J, Thompson K D, Morgan A L, et al. (2007) Seasonal variation and the immune response: A fish perspective. Fish & Shellfish Immunology 22(6): 695–706.

[59] Song Z, Zhang L Y, Dong H B, et al. (2012) Advances in JAK-STAT Signaling Pathway. China Animal Husbandry & Veterinary Medicine 39(06): 128–132.

[60] Pouliot P, Bergeron S, Marette A, et al. (2009) The role of protein tyrosine phosphatases in the regulation of allergic asthma: implication of TC-PTP and PTP-1B in the modulation of disease development. Immunology 128(4): 534–542.

[61] Morales J K, Falanga Y T, Depcrynski A, et al. (2010) Mast cell homeostasis and the JAK–STAT pathway. Genes & Immunity 11(8): 599–608.

[62] Ivashkiv L B, Hu X (2004) Signaling by STATs. Arthritis Res Ther 6(4): 159.

[63] Madhubanti Basu, Mahismita Paichha, Banikalyan Swain, et al. (2015) Modulation of TLR2, TLR4, TLR5, NOD1 and NOD2 receptor gene expressions and their downstream signaling molecules following thermal stress in the Indian major carp catla (*Catla catla*). 3 Biotech, 5(6):1021–1030.

[64] Bagnyukova T V, Lushchak O V, Storey K B, et al. (2007) Oxidative stress and antioxidant defense responses by goldfish tissues to acute change of temperature from 3 to 23°C. Journal of Thermal Biology 32(4): 227–234.

[65] Vergauwen L, Benoot D, Blust R, et al. (2010) Long-term warm or cold acclimation elicits a specific transcriptional response and affects energy metabolism in zebrafish. Comparative Biochemistry and Physiology - Part A: Molecular & Integrative Physiology 157(2): 149–157.

[66] Rawles S D, Green B W, Gaylord T G, et al. (2012) Response of sunshine bass (Morone chrysops x M. saxatilis) to digestible protein/dietary lipid density and ration size at summer culture temperatures in the Southern United States. Aquaculture 356-357: 80–90.

[67] Craig S R, Neill W H, Gatlin D M (1995) Effects of dietary lipid and environmental salinity on growth, body composition, and cold tolerance of juvenile red drum (*Sciaenops ocellatus*). Fish Physiology and Biochemistry 14(1): 49–61.

